# Atf3 controls transitioning in female mitochondrial cardiomyopathy as identified by single-cell transcriptomics

**DOI:** 10.1101/2023.09.11.557118

**Authors:** Tasneem Qaqorh, Yusuke Takahashi, Kentaro Otani, Issei Yazawa, Yuya Nishida, Yoshitaka Fujihara, Mizuki Honda, Shinya Oki, Yasuyuki Ohkawa, David R. Thorburn, Ann E. Frazier, Atsuhito Takeda, Yoshihiko Ikeda, Heima Sakaguchi, Takuya Watanabe, Norihide Fukushima, Yasumasa Tsukamoto, Naomasa Makita, Osamu Yamaguchi, Kei Murayama, Akira Ohtake, Yasushi Okazaki, Hijiri Inoue, Ken Matsuoka, Satoru Yamazaki, Seiji Takashima, Yasunori Shintani

## Abstract

Oxidative phosphorylation defects results in mitochondrial diseases, with cardiac involvement markedly impacting prognosis. However, the mechanisms underlying the transition from compensation to dysfunction in response to metabolic deficiency remain unclear, impeding the development of effective treatments. Here, we employed single-nucleus RNA sequencing (snRNA-seq) on hearts from mitochondrial cardiomyopathy (MCM) mice with cardiac-specific *Ndufs6* knockdown (FS6KD). Pseudotime trajectory analysis of cardiomyocytes from early stage of female FS6KD hearts revealed dynamic cellular state transitioning from compensation to severe compromise, coincided with transient upregulation of a critical transcription factor, *activating transcription factor 3* (*Atf3)*. Genetic ablation or adeno-associated virus-mediated *Atf3* knockdown in FS6KD mice effectively delayed cardiomyopathy progression in a female-specific manner. Notably, human MCM snRNA-seq revealed a similar transition, including the dynamic expression of *ATF3*. In conclusion, our findings highlight a fate-determining role of Atf3 in female MCM progression, providing a promising therapeutic candidate for the currently intractable disease.

## Introduction

Abnormalities in oxidative phosphorylation (OXPHOS) lead to metabolic insufficiencies resulting in systemic mitochondrial diseases (MDs). Congenital mitochondrial disorders can arise from mutations in mitochondrial DNA (mtDNA) or genomic DNA ^1^. As numerous gene products contribute to mitochondrial integrity and OXPHOS homeostasis, more than 400 causative genes have been identified ^2^. This genetic diversity contributes to the high prevalence and heterogeneity of MDs, impacting about 1 in 4000 individuals ^3,4^. Moreover, emerging evidence highlights tissue-specific ^5^ and even cell-specific ^6^ responses. It should be noted that there are compensatory mechanisms for dysfunctional or damaged mitochondria, including mtDNA repair, mitochondrial dynamics (fission and fusion), and increased mitochondrial biogenesis ^7,8^. However, insults exceeding cellular compensatory capacity drive dysfunction, leading to tissue failure and disease manifestation. Although the effects are clear, the mechanisms underlying the transition from compensation to dysfunction in disease progression remain unknown. These complexities, coupled with an incomplete understanding of molecular mechanisms of disease progression, have hindered therapeutic development, so MD remains intractable.

Previous studies have elucidated stress response pathways activated by mitochondrial dysfunction, including the mitochondrial unfolded protein response and integrated stress response (ISR^mt^) ^9^. ISR^mt^ triggers eukaryotic translation initiation factor 2 alpha phosphorylation through upstream cascades, suppressing global translation and selectively inducing downstream players, such as activating transcription factor 4 (Atf4), Atf5, DNA damage–inducible transcript 3, (CHOP), and fibroblast growth factor 21 (Fgf21) ^10–17^. Although ISR^mt^ is acknowledged as a critical modulator of MD progression, its role is context-or model-dependent. Recently, Ahola *et al.* reported that activation of Oma1–Dele1–Atf4 axis in Cox10-deficient mice, an OXPHOS deficiency model, enhanced glutathione metabolism, protecting against ferroptosis ^18^. Inhibition of ISR^mt^ through Oma1 or Dele1 deletion, or use of an ISR inhibitor exacerbated the cardiomyopathy phenotype, suggesting a protective mechanism for stress attenuation. Conversely, in helicase twinkle-deficient (Deletor) mice ^16,17^, which accumulates mtDNA deletions with age, the Fgf21–Atf4–one carbon metabolism axis plays a detrimental role in progressive myopathy. ISR^mt^ may primarily serve as a protective response for cellular recovery, yet prolonged ISR^mt^ could trigger detrimental effects. Even when ISR^mt^ plays a protective role, accelerated tissue dysfunction persists despite the inhibition of ISR^mt^-associated mediators and downstream players, suggesting the existence of an independent trigger for tissue dysfunction ^18,19^. Hence, identifying upstream factors or mechanisms driving the transition from compensation to dysfunction prior to ISR^mt^ activation is crucial. Moreover, understanding initial triggers is critical for establishing effective therapeutic strategies.

To gain insights into the regulatory mechanisms underlying MD progression, we employed single-nucleus RNA sequencing (snRNA-seq) on heart samples collected at different time points from mitochondrial cardiomyopathy (MCM) mice with cardiac-specific *Ndufs6* knockdown (FS6KD) ^20^. Pseudotime trajectory analysis of cardiomyocytes from the early-stage of female FS6KD cardiomyocytes, followed by differentially expressed gene (DEG) analysis, revealed dynamic transitioning from compensation to severely compromise, steered by a pivotal transcription factor (TF), *Atf3*. Notably, snRNA-seq also revealed that this dynamic transition was recapitulated in the human MCM hearts.

## Results

### Early compensatory upregulation of *Ppargc1a* precedes cardiomyocyte dysfunction

MCM, which is currently garnering research attention ^21^ can manifest independently, although it is often associated with organ dysfunction, and contributes to mortality ^22^. In the present study, we used FS6KD mice, an MCM model with heart-dominant knockdown of the complex I subunit in the mitochondrial respiratory chain, because decreased complex I activity is the predominant defect among patients with MD ^23–26^, and the model allows noninvasive cardiac function measurement. Unlike male mice, which exhibit a low left ventricular ejection fraction (EF) beginning at an early age and rapidly develop cardiac failure ^20^, female FS6KD mice showed a gradually progressing decline in left ventricular EF% beginning at 6–8 weeks (Figure S1A). We reasoned that a slowly progressing disease model would be suitable for dissecting transitions, thus, we used female FS6KD mice hereafter for further analysis. To comprehend disease development mechanisms and cell type-specific responses, we employed snRNA-seq on heart nuclei extracted from heart tissues of FS6WT mice and FS6KD mice at different disease stages: early-stage 8-week FS6KD and late-stage 17-week FS6KD.

Unsupervised clustering of integrated datasets revealed distinct cardiac cell populations, including cardiomyocytes, fibroblasts, immune cells, and minor populations, such as myoblasts (Figures 1A and S1B). Cellular composition variations were evident, with cardiomyocytes increasing and fibroblasts decreasing significantly in 8-week FS6KD hearts, and this change was inverted in 17-week FS6KD hearts (Figure S1C). Notably, both epicardial and endocardial populations decreased in FS6KD, whereas adipocytes were exclusive to the wild type (Figure S1D). Metabolic dysfunction prompted cellular state shifts, most prominently in cardiomyocytes, as observed in disease stage-specific clusters (Figure 1B). *Ndufs6* knockdown levels were similar across cell types (Figures 1C), indicating inherent cell-specific responses driving the observed shifts. Cardiomyocyte transcriptome assessed by DEG analysis revealed dynamic changes in metabolic regulation at 8 weeks, including increased fatty acid oxidation (FAO) and glycolytic shift, as reflected in significant gene ontologies (GOs). The genes marked at 8 weeks were downregulated at 17 weeks, whereas genes often associated with heart failure were upregulated at 17 weeks, with upregulation in genes enriched in response to muscle stretch GO terms (Figure 1D). This suggests that upregulation of processes in response to metabolic deficiency in 8-week cardiomyocytes failed to halt disease progression. Analysis of isolated and subclustered cardiomyocytes (Figures 1E and 1F) showed the most significant DEG in 8 weeks was peroxisome proliferator–activated receptor gamma coactivator 1-alpha (*Ppargc1a*) (Figures 1G and 1I), indicating its potential role in such adaptive responses. *Ppargc1a*, a transcription co-activator and mitochondrial biogenesis master regulator ^27^, adapts cells to increased energy needs. Conversely, heart failure marker natriuretic peptide B (*Nppb*) ^28^ was upregulated in the 17-week cardiomyocytes (Figure 1H).

**Figure 1.**
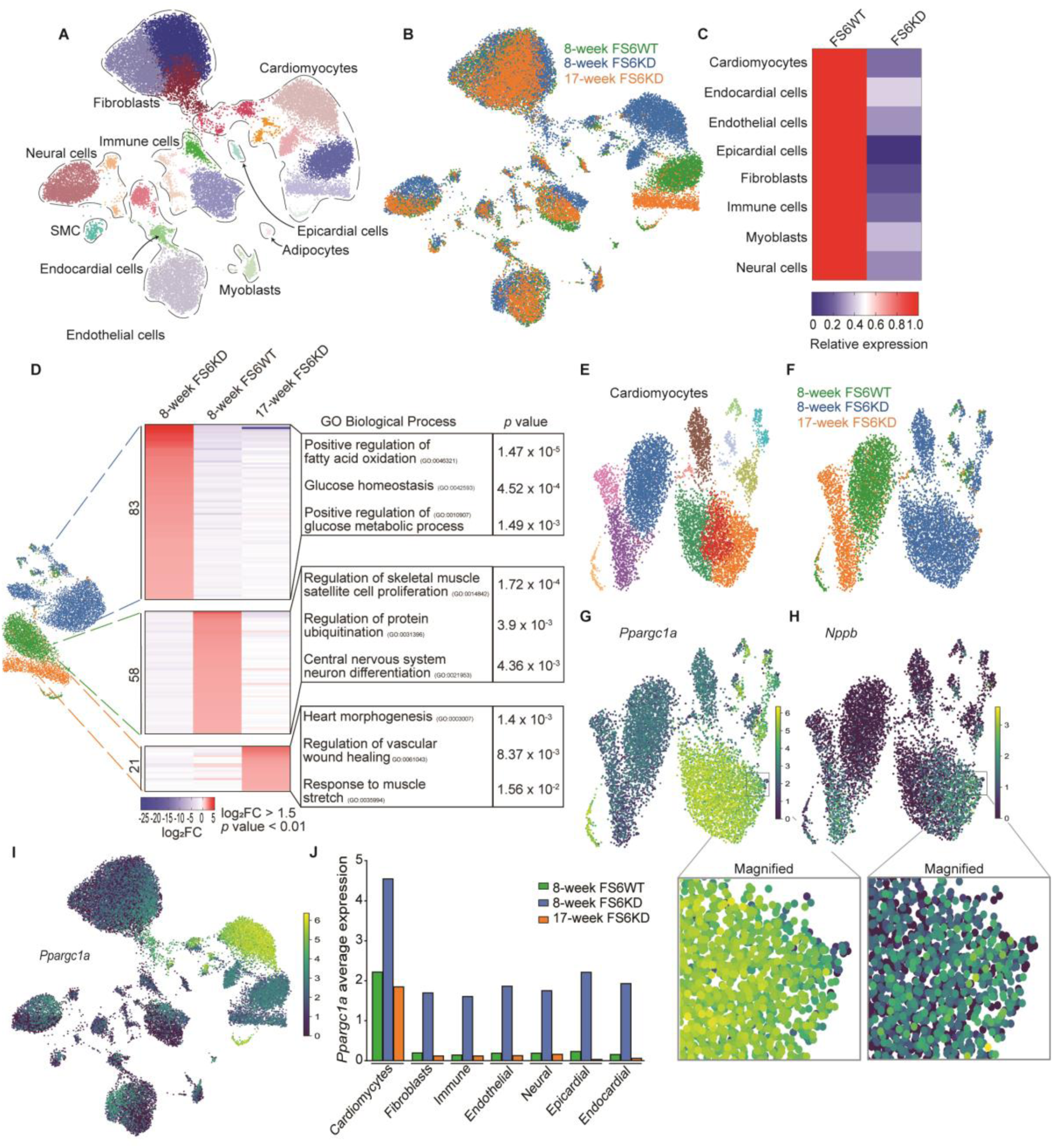
Upregulation of *Ppargc1a* in 8-week FS6KD cardiomyocytes for metabolic compensation. (A and B) Uniform manifold approximation and projection (UMAP) plot of the integrated dataset cardiac cellular populations. (A) Colored by leiden clusters and annotated. (B) Colored by sample identity, 8-week FS6WT (Green), 8-week FS6KD (Blue), and 17-week FS6KD (Orange), n = 1 per group. (C) Heatmap displaying knockdown efficiency per cell type. Relative average expression levels of *Ndufs6* in cell types of 8-week FS6KD against 8-week FS6WT. (D) UMAP plot of isolated cardiomyocytes colored by sample identity with respective heatmaps displaying top upregulated genes ranked by log_2_FC and associated enriched top GO terms. Fisher exact test was used to calculate *p*-values for GO. (E and F) UMAP plot of isolated and subclustered cardiomyocytes. (E) Colored by leiden clusters. (F) Colored by sample identity. (G) *Ppargc1a* expression shown as isolated cardiomyocytes UMAP feature. Small subcluster from 8-week FS6KD was magnified and shown in lower panel highlighting decreased expression. (H) *Nppb* expression shown as isolated cardiomyocytes UMAP feature. Small subcluster from 8-week FS6KD was magnified and shown in lower panel highlighting increased expression. (I) *Ppargc1a* expression shown as integrated dataset UMAP feature. Color intensity is increased in cellular populations corresponding to 8-week FS6KD in all cell types. (J) *Ppargc1a* average expression per cell type per dataset.

Despite the ongoing metabolic burden and its role in cardiac adaptation to energy depletion ^27^, *Ppargc1a* was markedly attenuated in 17-week cardiomyocyte cluster (Figures 1G and 1I). Notably, in a small subcluster of 8-week cardiomyocytes, *Ppargc1a* was downregulated with concurrent *Nppb* upregulation, resembling 17-week cardiomyocytes (Figures 1G and 1H; magnified lower panels). These findings suggest the presence of ongoing processes driving the transition of 8-week cardiomyocytes toward a maladaptive state, similar to that of 17-week cardiomyocytes. Other cardiac cell responses to metabolic deficiency showed consistent *Ppargc1a* upregulation at 8 weeks across all cell types, with pronounced elevation in cardiomyocytes (Figures 1I and 1J). Moreover, all cell types were restored to baseline levels by 17 weeks. Importantly, under normal conditions, cardiomyocytes exhibited significantly higher baseline *Ppargc1a* expression, indicating their heightened metabolic demands and the probable susceptibility to metabolic perturbations.

### Pseudotime trajectory analysis reveals *Atf3* induction during the transition

Given the heterogeneous subclusters found in 8-week FS6KD cardiomyocytes, especially the subcluster with downregulated *Ppargc1a* and upregulated *Nppb* (Figures 1G and 1H; magnified lower panels), resembling a maladaptive state, we hypothesize that a dynamic transition occurs from a compensatory to maladaptive across 8-week FS6KD cardiomyocytes. Employing pseudotime trajectory inference analysis ^29^, we identified a trajectory comprised of 8 consecutive cellular states (Figures 2A-2C). *Ppargc1a* expression consistently decreased through the trajectory to state 22 (Figure 2D) while *Nppb* expression increased in state 22 and 17-week FS6KD (Figure 2E). States 8 through 6 exhibited enrichment of metabolic regulatory processes switching to glycolytic metabolism (Figure 2F), akin to GOs found in 8-week FS6KD cardiomyocytes (Figure 1D). GO terms associated with reduced heart function were observed in states 10 and 22, corresponding to 17-week FS6KD cardiomyocytes ontology (Figures 2F and 1D). Thus, we labeled states 8 and 22 as early and late states, paralleling 8 and 17-week, respectively. Early-state cardiomyocytes (states 8 and 5) did not show upregulation of *Nppb* or other heart failure-related genes, similar to WT cardiomyocytes, but exhibited high *Ppargc1a* expression levels, suggesting compensation for mitochondrial dysfunction. Considering the similarity between the late states in 8-week trajectory and 17-week cardiomyocytes, we hypothesized that the cellular transition from compensation to dysfunction occurs in 8-week cardiomyocytes, with genes expressed during mid-trajectory states (states 4 through 12) influencing cellular fate transition.

**Figure 2.**
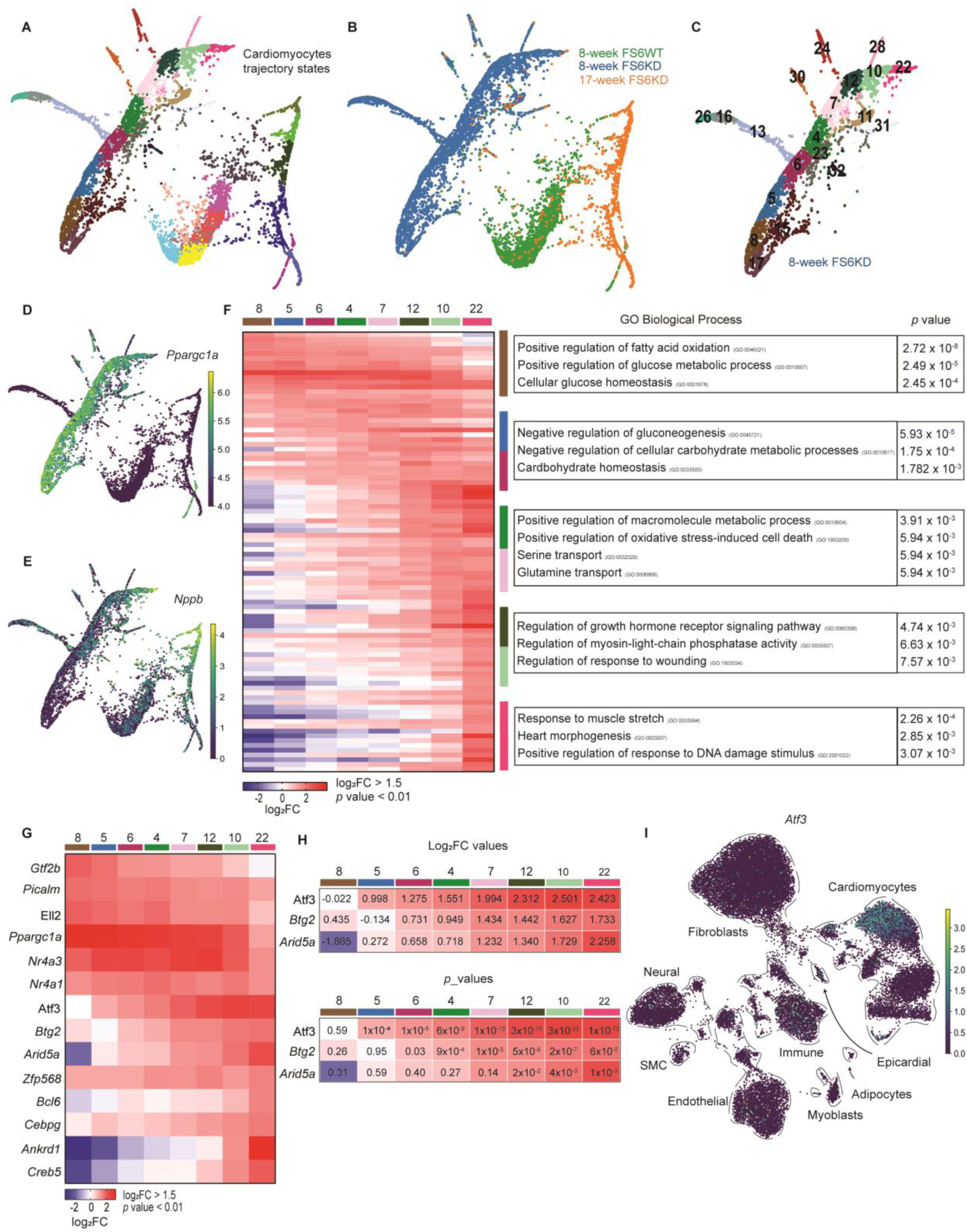
Pseudo-time trajectory analysis revealed *Atf3* induction during the transition. (A and B) ForceAtlas2 (FA2) scatter plot of abstracted partition-based graph abstraction (PAGA) trajectory from isolated cardiomyocytes. (A) Colored by leiden cellular states. (C) Colored by sample identity. (C) FA2 scatter plot of isolated PAGA trajectory from 8-week FS6KD cardiomyocytes numbered by leiden cellular states. (D) *Ppargc1a* expression shown as FA2 scatter plot. (E) *Nppb* expression shown as FA2 scatter plot. (F) Heatmap displaying top upregulated genes in the cellular states of 8-week FS6KD cardiomyocytes trajectory ranked by log_2_FC and associated top GO terms. Fisher exact test was used to calculate *p*-values for GO. (G) Heatmap displaying expression levels of transcription factors (TF) extracted from panel (F) ranked by log_2_FC. (H) Enlarged heatmap from panel (G) with log_2_FC values (upper panel) and *p*-values (lower panel). Wilcoxon rank-sum test was used to calculate *P*-values. (I) *Atf3* expression shown as integrated dataset UMAP feature. Color intensity is increased in 8-week FS6KD cardiomyocytes.

TFs play a crucial role in driving cell fate. We evaluated TF expression along the trajectory, summarizing it in a heatmap (Figure 2G). *Atf3* and *Btg2* are two TF genes whose expression was upregulated at states 4 and 7 with statistical significance, with Atf3 showing a higher fold change expression value and lower p-value (Figure 2H). Atf3 is a member of the ATF family that functions as a stress response factor. In contrast, *Btg2* is an immediate early gene induced by growth factors, and its function has been linked to cell cycle regulation or cardiac hypertrophy ^30,31^, a less likely candidate for transition factor in MCM. Moreover, *Atf3* was significantly upregulated in 8-week cardiomyocytes, downregulated in 17-week cardiomyocytes, and not expressed in other cell types or WT cardiomyocytes (Figure 2I). From these findings, we focused on Atf3, which might act as a transcriptional regulator, influencing cardiomyocyte fate toward dysfunction.

### Cluster of Atf3-expressing cardiomyocytes with repressed OXPHOS genes in the FS6KD heart

To validate our findings obtained from the trajectory analysis of snRNA-seq, we conducted a spatiotemporal assessment of the FS6KD heart. Immunostaining analysis of Atf3 in FS6KD cardiac tissues showed most of Atf3-expressing cells were cardiomyocytes, positive for the cardiomyocyte-specific marker TnnI3 (Figure 3A), consistent with the snRNA-seq results. Furthermore, Atf3 expression was restricted to young FS6KD mice (Figure 3B), not observed in WT and older FS6KD mice. Notably, clustered Atf3-positive signals were observed across cardiac sections, varying in intensity. Central nuclei exhibited higher signal intensity of the surroundings, indicating an expression gradient (Figure 3C). We hypothesized that *Ppargc1a* expression would be lower in nuclei with higher Atf3 levels and vice versa. Employing photo-isolation chemistry ^32,33^, an immunostaining-based method capable of extracting transcriptome information from locally defined areas by photo-irradiation, we separated the transcriptomes of cardiomyocytes with high or low Atf3 expression levels within Atf3-positive clusters. GO analysis of gene expression profiles in cells with high Atf3 expression levels revealed increased metabolic dysfunction, impaired respiration, and reduced cardiac contractility (Figure 3D). Remarkably, cells with high Atf3 expression displayed reduced *Ppargc1a* expression compared to cells with low Atf3 expression (Figure 3E). Overall, spatial transcriptomic profiles of contrasting Atf3-expressing cells corroborated our findings from the snRNA-seq trajectory analysis.

**Figure 3.**
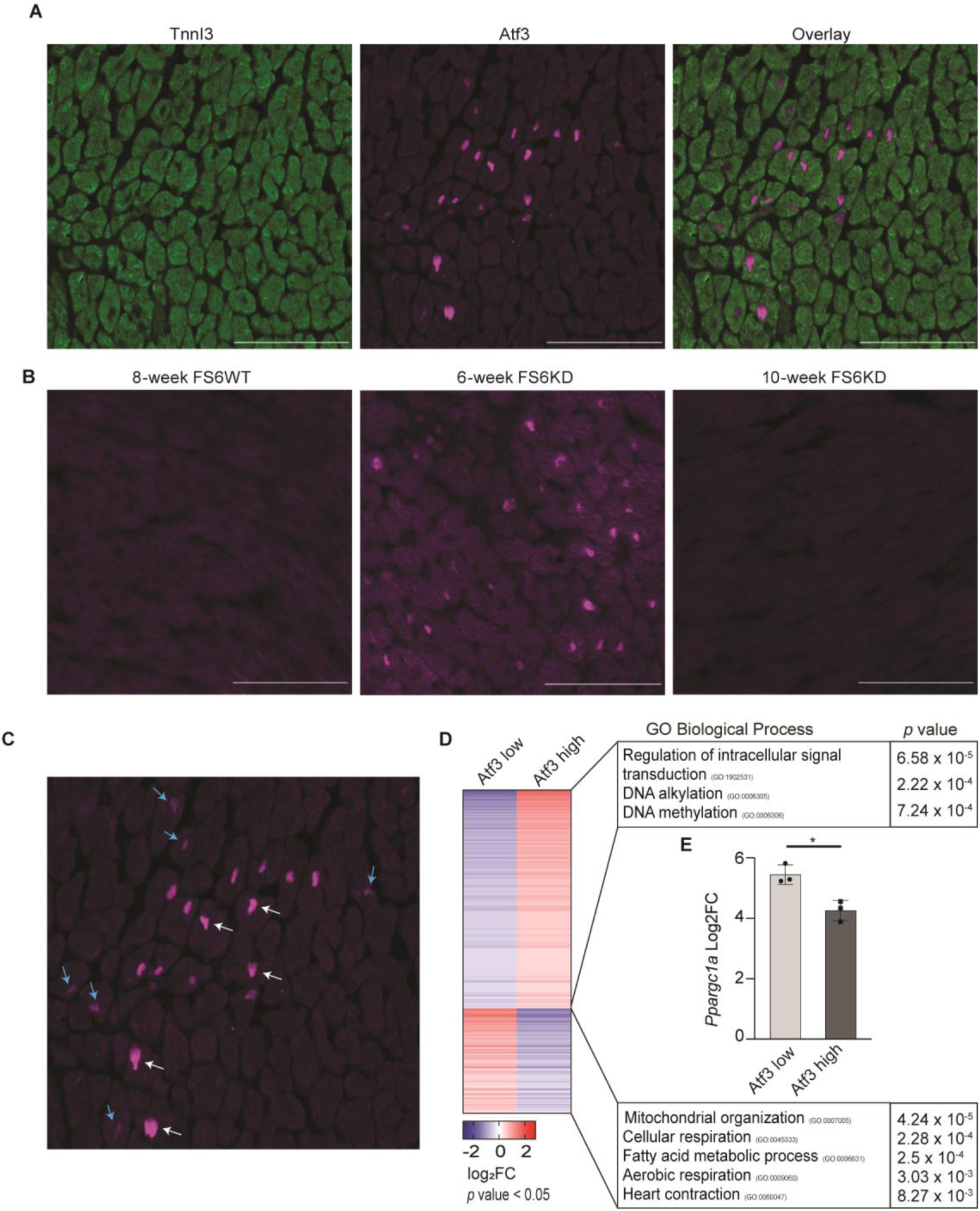
Atf3-expressing cardiomyocytes with repressed OXPHOS genes form cluster in the FS6KD heart. (A) Cardiomyocytes nuclear localization of Atf3 in 8-week FS6KD. Immunostaining with TnnI3 antibody (left), Atf3 antibody (middle), and an overlay image (right) (scale bar, 100 μm). (B) Expression and nuclear localization of Atf3 in FS6WT versus FS6KD of different ages. Immunostaining with Atf3 antibody in FS6WT (left), young 6-week FS6KD (middle), and old 10-week FS6KD (right) (scale bar, 100 μm). (C) Atf3 expression cluster from panel (A, middle). Blue arrows point to nuclei with low Atf3 expression and white arrows point to nuclei with high Atf3 expression (scale bar, 100 μm). (D) Heatmap displaying DEGs extracted from photo-isolation chemistry ranked by log_2_FC and associated top GO terms, n = 3 mice per group. Pairwise two-tailed t test was used to calculate *p*-values for DEG selection and Fisher exact test was used to calculate *p*-values for GO. (E) *Ppargc1a* expression levels. Bars: mean ± SD. Dots: individual subjects. Pairwise two-tailed t test, statistical significance: *p ≤ 0.05.

### ISR activation follows transient Atf3 induction in FS6KD cardiomyocytes

The ATF/CREB TF family have been extensively studied for their roles in ISR^mt^ upon mitochondrial dysfunction, primarily *Atf4* and *Atf5* ^10,17,18^. In contrast, *Atf3* has received less attention and is generally considered a minor downstream target of *Atf4* or *Atf5* ^16^. However, our investigations revealed no significant upregulation of *Atf4*, *Atf5*, or other ISR genes in 8 or 17-week FS6KD compared to WT. This observation led us to evaluate whether ISR^mt^ activation follows *Atf3* upregulation in the FS6KD heart. To this end, we introduced a 12-week snRNA-seq dataset as an intermediate state between 8 weeks and 17 weeks (Figures 4A and 4B). Cardiomyocytes exhibited the most pronounced changes, and clusters of 8-and 12-week cardiomyocytes were closely connected, implying that the 12-week dataset represented a state after the transition from 8 weeks. The transcriptome of cardiomyocytes at 12 weeks characterized by GOs included mitochondrial gene expression, mitochondrial translation, and pyruvate metabolism (Figure 4C), indicative of metabolic rewiring. These changes coincided with the upregulation of ISR response factors, such as *Atf4*, *Atf5*, and *Mthfd2* (Figures 4D-H). Importantly, expression of *Ppargc1a* and *Atf3* peaked at 8 weeks and returned to baseline at 12 weeks, supporting our hypothesis that ISR^mt^ activation followed *Ppargc1a* and *Atf3* downregulation. Moreover, 12-week cardiomyocytes exhibited induced expression of almost all subunits of mitochondrial respiratory chain complexes I and IV (Figure 4I), particularly assembly factors (*Ndufb10*, *Ndufb3*, *Cox6c*, *Cox7a2*). These might indicate a resisting attempt following the suppression of *Ppargc1a*, an upstream master regulator for mitochondrial biogenesis, at 8 weeks.

**Figure 4.**
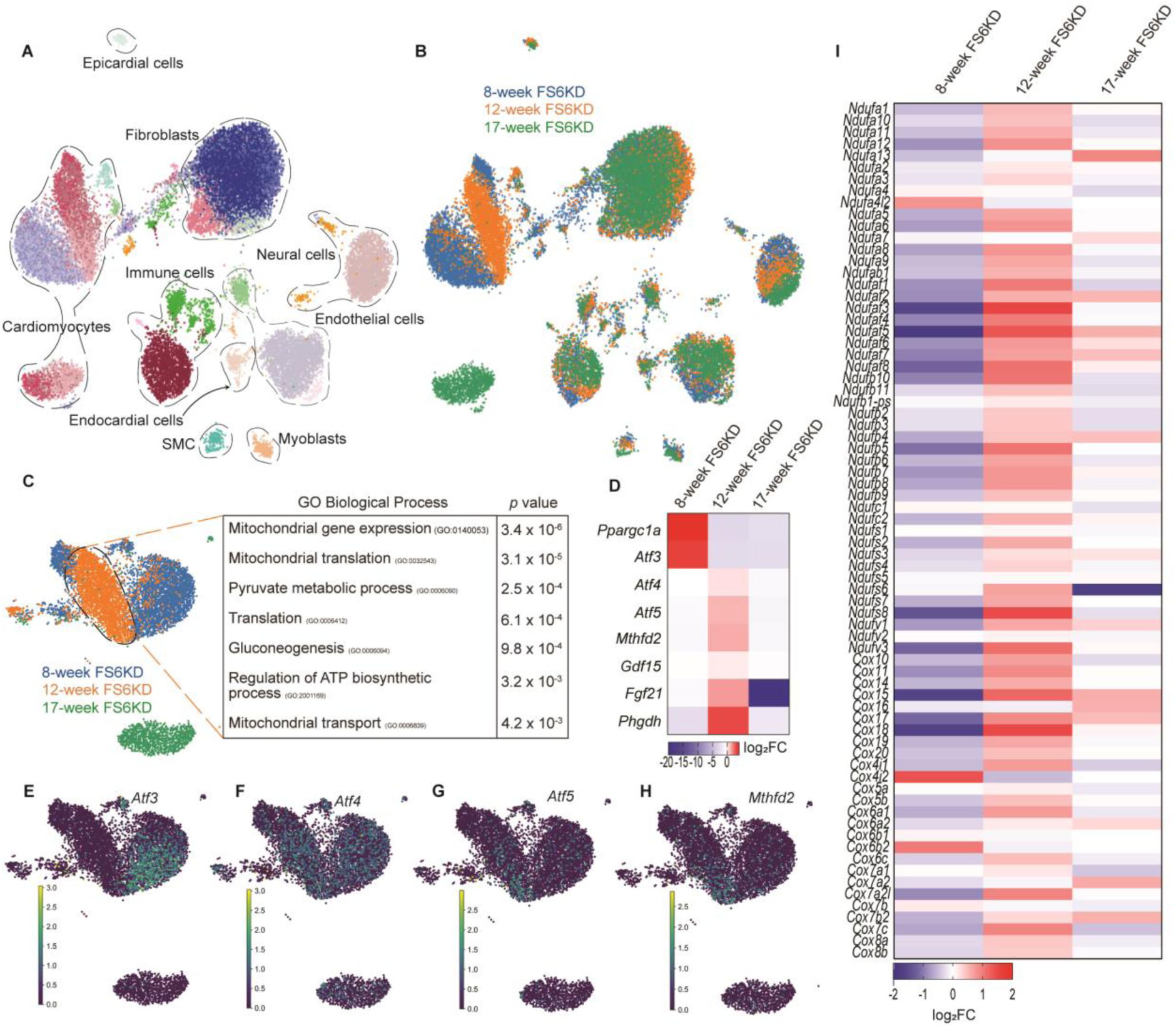
ISR activation follows transient *Atf3* induction in FS6KD cardiomyocytes. (A and B) UMAP plot of the integrated dataset cardiac cellular populations. (A) Colored by leiden clusters and annotated. (B) Colored by sample identity, 8-week FS6KD (Blue), 12-week FS6KD (Orange), and 17-week FS6KD (Green), n = 1 per group. (C) UMAP plot of isolated cardiomyocytes colored by sample identity and associated top GO terms of 12 weeks FS6KD cardiomyocytes. Fisher exact test was used to calculate *p*-values for GO. (D-H) Heatmap displaying expression levels of some *Atf*s and ISR^mt^ genes extracted from DEG list ranked by log_2_FC. (E-H) Expression shown as cardiomyocytes UMAP features of representative *Atf*s and ISR^mt^ genes (E) *Atf3*, (F) *Atf4*, (G) *Atf5*, and (H) *Mthfd2*. (I) Heatmap displaying expression levels of complex I and complex IV subunits and assembly factors ranked by log_2_FC.

### Genetic ablation and therapeutic knockdown of *Atf3* delays heart failure progression in female FS6KD mice

Our preceding findings highlight the role of *Atf3* expression in the transitioning cardiomyocytes from metabolic compensation to maladaptation during energy dysfunction. To assess whether the absence of *Atf3* improves cardiac function under OXPHOS dysfunction, we generated *Atf3*^−/−^FS6KD mice via a CRISPR/Cas9-mediated large deletion spanning exon 2 to 4 of *Atf3* in FS6KD fertilized eggs. Echocardiography analysis revealed a significantly improved EF at 8 weeks and demonstrated sustained stabilization of EF up to 12 weeks in *Atf3*^−/−^FS6KD, with a slight left ventricular diameter increase over time (Figures 5A and 5B). This suggests that *Atf3* is necessary to promote the decline in cardiac contractility observed in FS6KD mice and that *Atf3* knockout delayed heart failure progression in FS6KD mice. Notably, this cardioprotective effect disappeared in male counterparts (Figures 5C and 5D). Although we found the improved heart function in female *Atf3*^−/−^FS6KD, there was no significant decrease in fibrotic areas (Figures 5E-G). This suggests that cardiac fibrosis in FS6KD is likely not a response to decreased cardiomyocyte contraction but rather a reflection of fibroblast or other cell phenotypes due to mitochondrial dysfunction. *Atf3*^−/−^FS6WT mice displayed a normal cardiac phenotype, consistent with the previous report ^34^, implying that *Atf3* lacks a specific role under normal conditions and that our findings are not secondary to *Atf3* knockout effects. Next, we questioned if Atf3 may be leveraged as a therapeutic target. To determine whether cardiomyocyte-specific knockdown mimics whole-body knockout effects, we employed MyoAAV2-*shAtf3,* a muscle-directed adeno-associated virus (AAV capsid variant ^35^, to knock down *Atf3* in the muscles of neonatal FS6KD mice. After 8 weeks, echocardiography was conducted, and after 12 weeks, hearts were collected for RNA-seq using isolated heart tissue RNA (Figure 5H). MyoAAV2-sh*Atf3–*injected mice showed significantly higher EF compared with control (MyoAAV2-shCtrl) littermates (Figure 5I), reflecting the *Atf3*^−/−^FS6KD mice phenotype. This positive impact on cardiac function was gender-specific (Figure 5J), i.e., confined to females and absent in males.

**Figure 5.**
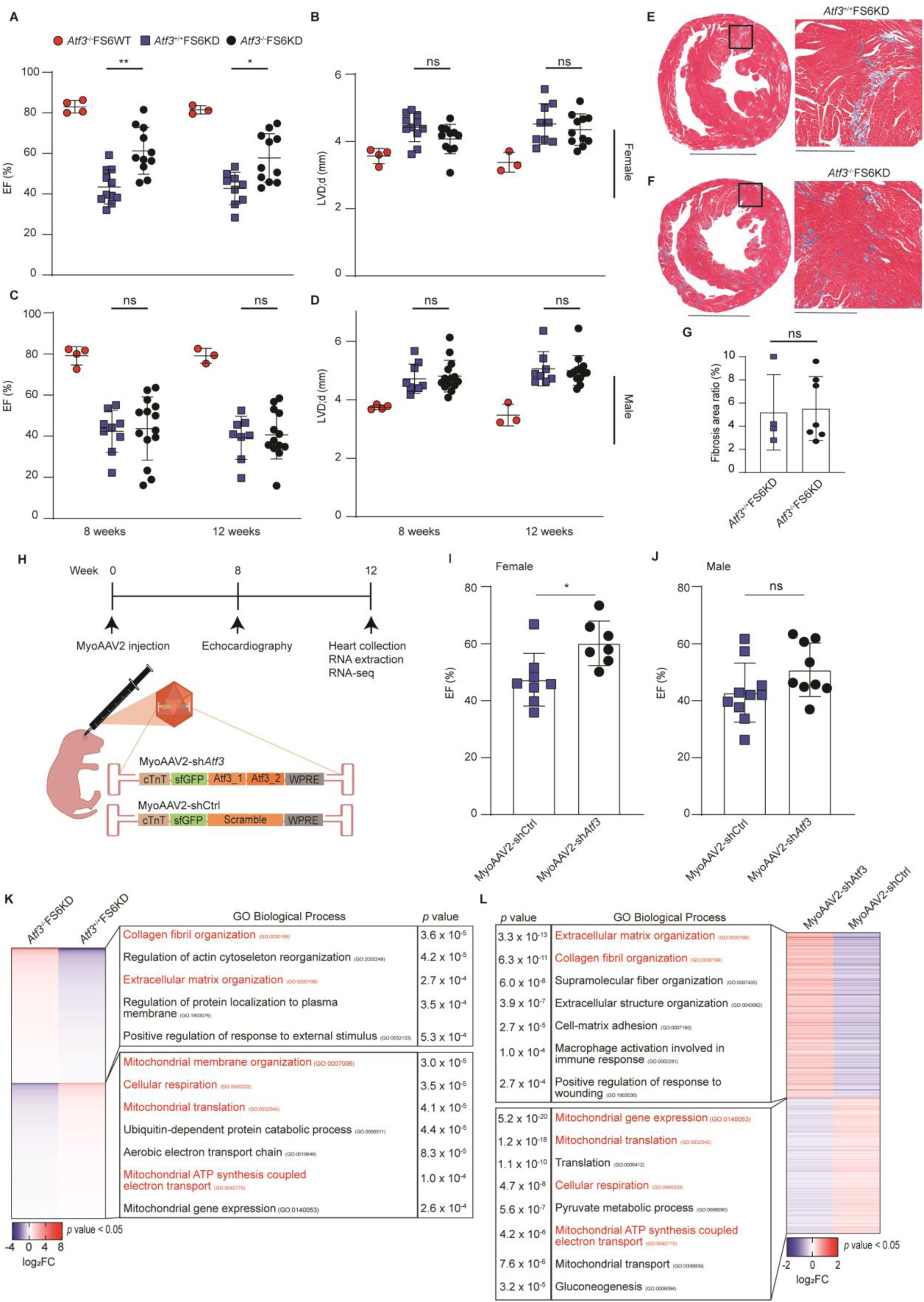
Genetic ablation of *Atf3* delayed the progression of heart failure in female FS6KD mice. (A-D) Summarized echocardiographic measurements of *Atf3*^-/-^FS6WT, *Atf3*^+/+^FS6KD, and *Atf3*^-/-^FS6KD mice at 8 weeks and 12 weeks. (A) EF and (B) LVDd of female mice, n = 4, 11, 11 for 8 weeks and 3, 9, 11 for 12 weeks. (C) EF and (D) LVDd of male mice, n = 4, 11, 11 for 8 weeks and 3, 9, 11 for 12 weeks, n = 4, 9, 14 for 8 weeks and 3, 8, 13 for 12 weeks. Bars: mean ± SD. Dots and squares: individual subjects. Two-way ANOVA and Tukey’s for multiple comparisons test, statistical significance: *p ≤ 0.05, **p ≤ 0.01, ns; not significant. (E-G) Masson’s Trichrome staining of hearts isolated from 12-week mice for fibrosis assessment. (E) Cross-section of *Atf3*^+/+^FS6KD (scale bar, 2.5 mm) with an enlarged microscopic view of the boxed areas in the left panel (scale bar, 500 μm). (F) Cross-section of *Atf3*^-/-^FS6KD (scale bar, 2.5 mm) with an enlarged microscopic view of the boxed areas in the left panel (scale bar, 500 μm). (G) Respective quantitative assessment summary of cardiac fibrosis area, n = 4 and 7. Bars: mean ± SD. Dots and squares: individual subjects. Unpaired two-tailed Student’s t test, statistical significance: ns; not significant. (H) Schematic graph of MyoAAV2-mediated knockdown workflow. (I-J) Comparison of EF of 8-week FS6KD mice injected with MyoAAV2-shCtrl or MyoAAV2-sh*Atf3*. (I) Female mice, n = 8 and 7. (J) Male mice, n = 10 and 9. Bars: mean ± SD. Dots and squares: individual subjects. Unpaired two-tailed Student’s t test, statistical significance: *p ≤ 0.05, ns; not significant. (K) Heatmap displaying DEGs extracted from RNA-seq of *Atf3*^+/+^FS6KD or *Atf3*^-/-^FS6KD hearts at 12 weeks ranked by log_2_FC and associated top GO terms, n = 4 mice per group. GO terms in red indicate similar terms found in panel (L). Pairwise two-tailed t test was used to calculate *p*-values for DEG selection and Fisher exact test was used to calculate *p*-values for GO. (L) Heatmap displaying DEGs extracted from RNA-seq of MyoAAV2-shCtrl or MyoAAV2-sh*Atf3*hearts at 12 weeks ranked by log_2_FC and associated top GO terms, n = 3 mice per group. GO terms in red indicate similar terms found in panel (K). Pairwise two-tailed t test was used to calculate *p*-values for DEG selection and Fisher exact test was used to calculate *p*-values for GO.

Notably, RNA-seq revealed that hearts of 12-week *Atf3*^−/−^FS6WT and MyoAAV2-*shAtf3* injected mice commonly exhibited downregulation of glycolytic and OXPHOS-dependent metabolic processes, as well as a reduction in mitochondrial maintenance such as translation and transport (Figures 5K and 5L). These GOs were oppositely upregulated in 12-week FS6KD cardiomyocytes (Figure 4C), presumably as a feedback mechanism after *Ppargc1a* and *Atf3* downregulation. This suggests that the loss of *Atf3* prevents cardiomyocytes from transitioning to a compromised state, in which downstream metabolic process genes were upregulated due to the loss of compensatory mitochondrial biogenesis orchestrated by *Ppargc1a*. From these observations, we conclude that *Atf3* plays a pivotal role in disease transitioning in female FS6KD mice and is a therapeutic candidate for female MCM.

### Cardiomyocytes of a human patient with complex IV deficiency display transitions

We aimed to determine whether our observations found in FS6KD mice were conserved in humans. However, due to the limited availability of heart tissue samples and their heterogeneity, as well as the transient nature of *Atf3* expression in young mice, capturing human patient expression patterns posed a challenge. Nonetheless, we conducted snRNA-seq on a small ventricular tissue sample from a 9-month-old female patient with MCM who exhibited severe complex IV reductions indicated by COX4 staining and characteristic abnormal cristae observed via electron microscopy (Figures 6A and 6B). Cardiac tissues were collected during ventricular assist device implantation as the patient suffered severe cardiac failure. Unsupervised clustering revealed multiple cellular populations, with cardiomyocytes comprising the largest (Figure 6C). Surprisingly, the expression of *ATF3*, *PPARGC1A*, and *NPPB* was observed in isolated cardiomyocytes (Figures 6D-6G). For validation, we integrated this data with data from healthy cardiomyocytes of an 11-year-old female donor heart ^36^. The healthy cardiomyocytes formed separate clusters, and did not exhibit upregulation of these genes (Figures S2A-S2D), confirming it was pathology-induced rather than an inherent higher human baseline. The presence of *ATF3* expression suggested the possibility of reconstructing a trajectory in human cardiomyocytes, prompting us to perform trajectory analysis. We identified two distinct paths (Figure 6H), but focused on the first trajectory, as it resembled gene expression patterns observed in 8-week FS6KD mouse cardiomyocytes. Specifically, we analyzed state 27, annotated as an early state based on high *PPARGC1A* expression (Figure S2E), through state 26, which showed significant *NPPB* upregulation (Figure S2F). A heatmap showed a gradual expression increase of *NPPB*, a clinically used biomarker for heart failure, along the trajectory, validating our trajectory analysis (Figure 6I).

**Figure 6.**
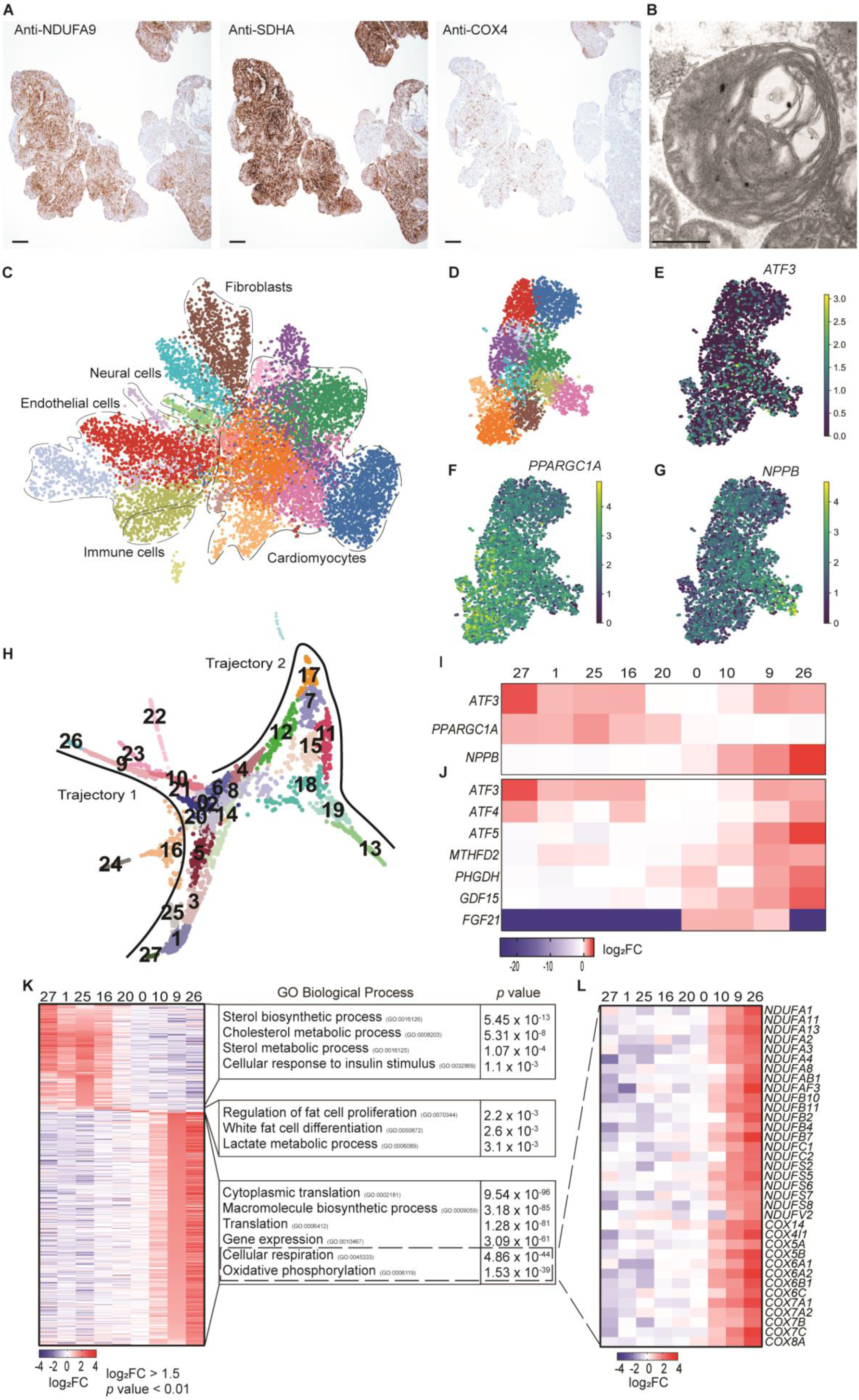
Cardiomyocytes of a patient with MCM demonstrated transitioning. (A) Immunohistochemical assessment of respiratory chain complexes abundance on myocardium from the female 9-month MCM patient. Immunostaining with NDUFA9 antibody (complex I, left), SDHA antibody (complex II, middle), and COX4 antibody (Complex IV, right) (scale bar, 200 μm). (B) Electron microscopy ultrastructure of abnormal cardiac mitochondria showing swelling and abnormal onion-like concentric membranes (scale bar, 500 nm). (C) UMAP plot of cardiac cellular populations from the MD patient colored by leiden clusters and annotated, n = 1. (D-G) UMAP plot of isolated cardiomyocytes (D) colored by leiden clusters. Expression shown as cardiomyocytes UMAP features of (E) *ATF3*, (F) *PPARGC1A*, and (G) *NPPB*. (H) FA2 scatter plot of PAGA trajectory from cardiomyocytes numbered by leiden cellular states, showing two trajectories. (I and J) Heatmap displaying expression levels of genes along the first trajectory states showing (I) *ATF3* expression level changes in comparison *PPARGC1A, NPPB* and (J) *ATF3* expression level changes in comparison to some ISR^mt^ genes ranked by log_2_FC. (K-L) Heatmap displaying top upregulated genes in the cellular states of (K) first trajectory selected cellular states ranked by log_2_FC and associated top GO terms. Fisher exact test was used to calculate *p*-values for GO. (L) Heatmap displaying expression levels of complex I and complex IV subunits and assembly factors extracted from the trajectory and ranked by log_2_FC.

DEG analysis of first trajectory highlighted upregulation of *ATF3* in the earliest state (state 27) and *PPARGC1A* expression in the initial states (states 27 to 25, Figure 6I), probably mirroring the late-trajectory states of 8-week FS6KD mouse (states 10 and 22 in Figure 2G). ATF family and ISR gene upregulation emerged in later states (Figure 6J). Given that 12-week mice expressed ISRs with suppressed *Atf3* levels (Figures 4D-4H), we hypothesized that the human cardiomyocyte cluster represents a more advanced phenotype than 8-week FS6KD mouse, extending the late-trajectory states of 8-week FS6KD mouse trajectory to 12-week FS6KD mice. Enrichment analysis of early human states revealed metabolic regulation processes, whereas late states showed enrichment of processes related to translation and upregulated gene expression (Figure 6K). Late states featured enriched aerobic respiration pathways and energy production, particularly in states 9 and 26, with complex I and IV subunits and assembly factors highly expressed (Figure 6L). Late state gene expression patterns resembled those of 12-week FS6KD mice (Figure 4C), strengthening the notion that the human trajectory extends from late 8-week FS6KD mouse states. In contrast, the DEGs in the second trajectory did not highlight heart failure-related genes. GO enrichment analysis revealed variations in metabolic regulation across all states (Figures S2G and S2H). Second trajectory may provide insights into different aspects of how mouse and human systems respond to mitochondrial deficiency or difference in the responsible damaged respiratory chain complex for MCM. Overall, our snRNA-seq from human MCM confirmed that the cardiomyocytes in severe heart failure still hold heterogeneity and operate dynamic transitioning. Notably, the *ATF3*-involving trajectory is likely preserved across species.

## Discussion

Understanding the initial mechanisms underlying the progression of MDs toward dysfunction is necessary to develop effective therapeutic strategies. Our snRNA-seq revealed dynamic transcriptional changes occurring during disease progression in the FS6KD heart. Notably, cardiomyocyte-specific transient upregulation of *Atf3* preceded ISR^mt^ and functioned as a pivotal trigger for transitioning from a compensatory state to maladaptation, facilitating the onset of female MCM. Importantly, snRNA-seq analysis of human MCM with complex IV deficiency indicated the existence of a trajectory marked by *ATF3* expression within cardiomyocytes. This finding shows that human cardiomyocytes undergo a similar process of compensation, ISR^mt^ activation, and maladaptation. Our findings therefore unveil an upstream stress-modulating axis in female MCM preserved across species.

Our data revealed that ISR^mt^ expression, followed not only *Atf3* induction, but also its subsequent downregulation. These finding aligns with our initial hypothesis that the ISR^mt^ genetic program is not the earliest response in MDs. Other models have demonstrated a paralleled upregulation of ISR^mt^ with severe myopathy phenotype. For instance, Oma1 activation in Cox10-deficient mice was observed simultaneously with severe skeletal muscle mass reduction and cardiac enlargement ^18^. Similarly, in Deletor mice with cumulative mtDNA deletions ^16^, although representing a milder OXPHOS deficiency model, the initial signs of ISR^mt^ coincided with respiratory chain deficiency. Kim, et al. reported that induction of *Atf3* plays a pivotal role in disease progression in the livers of Zucker diabetic fatty (ZDF) rats, a model of type 2 diabetes exhibiting hepatic steatosis, and patients with non-alcoholic fatty liver disease ^37^. ZDF rats demonstrate the induction of metabolic stress, impaired mitochondrial function, endoplasmic reticulum (ER) stress, and substantial reductions in fatty acid oxidation (FAO). Silencing *Atf3* resulted in the re-elevation of *Ppargc1a* that was downregulated along with disease progression, thereby restoring FAO levels and mitochondrial function, counterbalancing metabolic dysfunction as well as redox state. Notably, even when ER stress is evident, patients with low ATF3 levels exhibited diminished CHOP expression ^37^, suggesting the potential of ATF3 as a valuable biomarker. As an acute stress response factor, it is not surprising Atf3 induction is evident in various acute injury models ^34,38–43^. However, Atf3 likely plays a distinct role in chronic metabolic dysfunction models, such as MCM in the present study and ZDF rats. Capturing early Atf3 induction in MD models is challenging owing to their rapid progression and early-onset manifestations across multiple organs. Elucidating the upstream factors governing Atf3 fate-determination in MCM, particularly in FS6KD or similar models, necessitates further investigation.

A particularly intriguing finding was the gender-specific protective effect of Atf3 loss in FS6KD mice, observed exclusively in female mice and absent in males. Female mice were chosen owing to their slower disease progression, enabling the tracking of early disease-related responses. However, this protective response was not observed in male mice, contrary to expectations. Gender disparities manifest in various diseases, particularly cardiovascular ^44,45^, autoimmune ^46^, Parkinson’s, and Alzheimer’s diseases ^47^. Over the past few decades, studies have also shown gender disparities in mitochondrial function ^48^. In addition, a meta-analysis illustrated consistent gender differences in two domains of mitochondrial biology: higher mitochondrial content in women and elevated ROS production in men ^49^. This difference is most likely attributed to sex-hormones. For instance, the estrogen-related receptor family is involved in mitochondrial biogenesis and metabolic gene expression ^48^. Estrogen receptors are distributed across mitochondria, nuclei, and plasma membranes, and estrogen’s interaction with mtDNA affects transcription and replication, boosting mitochondrial-related gene expression ^50^. Female mice display enhanced fatty acids utilization and upregulate genes associated with FAO in muscles during intense exercise to a greater extent than males ^51^. Estrogen treatment enhances mitochondrial biogenesis, ATP production, and Ppargc1a expression in hearts after trauma-hemorrage ^52^. Ppargc1a coactivates estrogen receptor-alpha expression ^53^, and its absence increases susceptibility to dilated heart failure in female mice ^54^, underscoring the role of female sex hormones and their regulation of Ppargc1a in adapting to compromised metabolism. Although sex differences exist, it is possible that male mice experience faster transitions leading to Atf3 upregulation, with any protective effects dissipating before the 8-week cardiac assessment.

The most remarkable finding is the striking resemblance of the FS6KD phenotype and the cardiomyocytes trajectory from a patient with MCM. Despite the patient’s severe cardiac failure requiring ventricular assist device implantation, observing an ongoing transition process within the cells suggests a broader therapeutic intervention window than initially conceived. Interestingly, human cardiomyocytes displayed an additional trajectory absent in FS6KD mice, emphasizing the intricate heterogeneity of human tissues and the multifaceted stress responses characteristic of patients with MD. This heterogeneity could have stemmed from the respiratory chain complex-specific phenotype due to complex IV deficiency, distinct from complex I deficiency in FS6KD mice. Nonetheless, the presence of Atf3-mediated cellular dysfunction amidst such complexity underscores its significance, surpassing species boundaries.

In conclusion, understanding the early mechanisms driving MDs toward dysfunction is pivotal for developing effective treatments. This study reveals that Atf3 plays a crucial role in this process, orchestrating the shift of cardiomyocytes from compensation to maladaptation. Although further research is essential, these findings offer a novel insight into how stress responses and adaptations operate in MDs. Moreover, the parallels observed in patient-derived cardiomyocytes highlight the potential therapeutic benefits of targeting pathways involving Atf3 in MCM.

### Limitations of Study

*Atf3* upregulation initiates the shift of cardiomyocytes from a compensated metabolic state to a severe phenotype in both FS6KD mice and a patient with MCM. To validate this consistency across diverse MCM or MD manifestations in humans, tracking the phenotype over time is crucial. However, obtaining successive biopsies is challenging owing to the invasive nature of the procedure. Although ongoing cellular adaptation was observed, early onset disease manifestation hinders precise prediction of the timing for the *Atf3* surge. This temporal uncertainty complicates identifying the optimal therapeutic intervention window.

## Acknowledgments

First, we thank Dr. Mohammadsharif Tabebordbar at MIT, Dr. Ko Toshiyuki, and Dr. Seitaro Nomura at Tokyo University for helping us perform the MyoAAV2 experiment as a collaboration. All authors greatly thank Ms. Akiko Ogai, Ms. Satomi Kobayashi, Ms. Kayoko Ohashi, and Ms. Sonoko Adachi for laboratory assistance; laboratory members of Molecular Pharmacology for help and discussion; Dr. Daisuke Okuzaki, Dr. Daisuke Motooka and members of Genome Information Research Center, Research Institute for Microbial Disease, Osaka University for the help in data acquisition using the next-generation sequencer; Dr. Keisuke Nimura for thoughtful advice on snRNA-seq. This work was supported by funding from AMED under Grant Number JP21ek0109499 to Y.S., Grants-in-Aid for Scientific Research from the Japan Society for the Promotion of Science under Grant Number 23K18295 to Y.S.; 23H00372, 22H04676, and 22K19275 to Y.O., an Intramural Research Fund for Cardiovascular Diseases of National Cerebral and Cardiovascular Center under Grant Number 21-2-7 to Y.S., research grants from the Medical Research Center Initiative for High Depth Omics/the Cooperative Research Project Program of the Medical Institute of Bioregulation at Kyushu University to Y.O., and research grants from Miyata foundation bounty for Pediatric cardiovascular research and SENSHIN medical research foundation to Y.S. D.R.T. and A.E.F. were funded by grants from the NHMRC and the Mito Foundation, with work performed at the Murdoch Children’s Research Institute supported by the Victorian Government’s Operational Infrastructure Support Program. We thank Enago for English language editing.

## Authors contribution

T.Q. prepared samples, curated and managed data, conducted all sequencing computational analysis, and interpreted data. T.Q. and Y.S. performed study investigation; interpreted data, designed validation experiments and analysis, and prepared figures. Y.T. conducted FACS and singulator nuclei collection. I.Y. collected animal samples, contributed to RNA-seq and data analysis. Y.N. contributed to computational tools and environment setup. Y.T., I.Y., and Y.N. participated in data interpretation and figure preparation. K.O. conducted AAV knockdown experiments. Y.F. conducted CRISPR/Cas9 knockout production. M.H., S.O., and Y.O. conducted PIC experiment, data processing and participated in data interpretation. D.R.T. and A.E.F. provided the animal model. A.T., K.M., A.O., and Y.I conducted diagnosis of human samples with A.T. conducting tissue immunohistology and Y.I. conducting EM analysis. H.S., N.M., O.Y., T.W., N.F., and Y.T. collected and provided human samples. H.I., K.M., S.Y., and S.T provided critical review, discussion, and feedback. Y.S. conceptualized the study, experimental design, secured funding, and provided overall supervision. T.Q. and Y.S. wrote the manuscript, which was edited and approved by all co-authors.

## Methods

### Animal models

*Ndufs6*^gt/gt^ heterozygous mouse sperm of mixed genetic background, C57BL/6J and 129/Ola, was kindly provided by Dr. David Thorburn ^20^. The mice were housed in a specific pathogen-free animal facility with a 12-hour light cycle and given a regular chow diet and all experiments involving animals followed the guidelines of National Cerebral and Cardiovascular Center. *Atf3*^-/-^FS6KD mice were created using CRISPR/Cas9 system as described previously^55^ by introducing gRNA/CAS9 (5’-ccagcgcagaggacatccga-3’ for the exon 2 of *Atf3* and 5’-cccagcagccaagagccgtt-3’ for the exon 4 of *Atf3*) protein solution into fertilized eggs of *Ndufs6*^gt/gt^ mice with an electroporator (NEPA21, Nepagene, Chiba, Japan). We obtained F0 founders that had a large deletion in the coding regions of *Atf3*, then intercrossed them to produce functional Atf3-null mice with FS6KD background and Atf3-null mice without FS6*^gt^* allele. The genotyping primers used were: 5’-gaggtaggctgtcagaccccatgc-3’ for Pr.1, 5’-gcccattctcgggtgcacactatacc-3’ for Pr.2, and 5’-gccacagtggaggatgtggtccc-3’ for Pr.3. Mice with muscle-specific knockdown were created by injecting *Ndufs6*^gt/gt^ neonatal mice with 50 µl MyoAAV2 containing 2 x 10^11^ GC, diluted with 0.001% Poloxamer 188 (Sigma) in PBS targeting mouse *Atf3* (pAAV[2miR30]-cTnT>sfGFP:{GAGCCTGGTGTTGTGCTATTTA}:{GAGATTCGCCATCCAGAATAAA}: WPRE). As a negative control, mice were injected with similar concentrations of scramble myoAAV2 (pAAV[miR30]-cTnT>sfGFP:Scramble[ACCTAAGGTTAAGTCGCCCTCG]:WPRE). MyoAAV2s used in this study were constructed and packaged by VectorBuilder and were administered by retro-orbital vein injections. All the different genotypes were born in the expected Mendelian ratios and were viable. All experiments and analyses were conducted using female mice.

### Echocardiography

Prior to echocardiography, mice were anesthetized and maintained with 0.25% isoflurane administered via a mask covering the nose and mouth of the animals. Transthoracic echocardiography was performed on unconscious mice with a Vevo 3100 imaging system (Visualsonics, Inc.). M-mode echocardiographic images were obtained from a short-axis view to measure the left ventricular ejection fraction (EF).

### Single-nuclei transcriptomics

#### Nuclei isolation from mouse heart tissues

Nuclei were isolated according to Frankenstein protocol ^56^ with minor modifications. All samples, reagents and steps were kept and performed on ice. 80 mg heart tissue was cut into small pieces in 0.5 ml ice-cold EZ PREP lysis buffer (Sigma) and homogenized using 2 ml glass douncer (8 times with pestel A and 1 time with pestel B) and incubated on ice with additional 1 ml ice-cold EZ PREP lysis buffer for 3 min with gentle mixing using a wide bore tip two times. Homogenate was filtered through a 70 μm followed by 20 μm cell strainer (PluriSelect) and centrifuged at 500 x g for 5 min at 4°C to remove cell debris and aggregates. Nuclei pellet was suspended in another 1.5 ml ice-cold EZ PREP lysis buffer and incubated on ice for 3 min. After centrifugation at 500 x g for 5 min at 4°C, nuclei pellet was incubated with nuclei washing and resuspension buffer consisting of 1X PBS, 1% BSA, 0.1% Tween-20 and 0.2 U/μl RNase inhibitor (TAKARA Bio) for 5 min before resuspension with additional 1 ml. Nuclei were centrifuged at 500 x g for 5 min at 4°C and the nuclei pellet was resuspended with 0.5 ml nuclei washing and resuspension buffer containing DAPI stain (DOJINDO) for nuclei labeling. Final nuclei count was determined by counting nuclei with a hemocytometer (NanoEntek) by fluorescent microscopy (Keyence BZ-X810).

#### FACS sorting

Isolated nuclei, kept on ice, were immediately sorted by Fluorescence Activated Cell Sorting using a FACS Aria Fusion cell sorter (Becton Dickinson, National Cerebral and Cardiovascular center). The 355 nm ultraviolet laser was used for analysis and nuclei sorting, while scattering detection of 488 nm blue laser was used to record forward-scatter characteristics (FSC) and side-scatter characteristics (SSC). Data were recorded and analyzed for gating and single nuclei selection using BD FACS DIVA (ver. 8.0.3) software. Final samples were centrifuged at 500 x g for 10 min at 4°C and pellets were resuspended with nuclei washing and resuspension buffer not containing Tween-20, to a final concentration of 1000 nuclei/μl. Final nuclei count was determined by counting nuclei with a hemocytometer (NanoEntek) using an all-in-one microscope (Keyence, BZ-X810).

### Human mitochondrial cardiomyopathy heart tissues

The procedure for snRNA-seq of heart tissues isolated from a patient with MCM was approved by the ethics committee of National Cerebral and Cardiovascular Center (Approval No. R20035). The cardiac tissues were obtained during ventricular assist device implantation, and frozen at −80°C immersed in Bambanker solution (NIPPON Genetics) until analysis. Diagnosis of MCM was based on severe reductions in CIV assessed by immunohistochemistry ^57^, enzyme activity ^58^, and characteristic mitochondrial features assessed by electron microscopy.

### Nuclei isolation from human heart tissues

Nuclei were isolated using Singulator 100 system (S2 Genomics, Inc) according to manufacturer’s recommendations. All samples, reagents and steps were kept and performed on ice. Small 11 mg heart tissue was cut into small pieces and inserted into nuclei cartridge (S2 Genomics, Inc) with 0.2 U/μl RNase inhibitor. Default protocol 2a was selected with one disruption cycle and 5 min incubation post homogenization in 4 ml Nuclei Isolation Reagent (S2 Genomics, Inc). Sample was passed through 40 μm in the cartridge, followed by a rinse with Nuclei Storage Reagent (S2 Genomics, Inc). Collected nuclei were centrifuged at 500 x g for 10 min at 4°C and pellets were resuspended with 4 ml nuclei washing and resuspension buffer containing 1X PBS, 1% BSA, and 0.2 U/μl RNase inhibitor. Additional centrifugation at 500 x g for 5 min at 4°C was performed and the nuclei pellet was resuspended with washing and resuspension buffer containing DAPI stain (DOJINDO) for nuclei labeling. Final nuclei count was determined by counting nuclei with a hemocytometer (NanoEntek) by fluorescent microscopy (Keyence BZ-X810).

### Library preparation, sequencing, and reads processing

Droplets capturing single nucleus from whole heart nuclei suspension were used for library generation in the 10X Genomics Chromium controller according to the manufacturer’s instructions in the Chromium Single Cell 3′ Reagent Kit v.3 User Guide. Additional components used for library preparation include the Chromium Single Cell 3′ Library and Gel Bead Kit v.3 (PN-1000092). Libraries were prepared according to the manufacturer’s instructions using the Chromium Single Cell 3′ Library and Gel Bead Kit v.3 (PN-1000092) and Chromium i7 Multiplex Kit (PN-120262). Final libraries were sequenced on Illumina NovaSeq6000. All libraries were sequenced to a depth of at least 20,000 total mean reads per nucleus. Cell Ranger (ver. 5.0.0 for mouse and ver. 6.0.0 for human) pipeline provided by 10X Genomics was used to process raw sequencing reads. Reads were aligned to mouse mm10 and to human GRCh38 references with pre-mRNA intronic regions. Gene-barcode matrices were created by quantifying UMI counts per gene per cell and the data were aggregated and normalized creating a combined gene-barcode matrix. Individual matrices of samples were used for downstream processing and analysis.

### Mouse samples integration, gene matrix pre-processing, filtering, and cell clustering

Python’s (ver. 3.8.5) Scanpy ^59^ (ver. 1.8.2) package was used for further pre-processing, filtering, and clustering of gene matrices obtained by Cell Ranger. For comparison between two or more datasets, ComBat ^60^ algorithm was used for batch effect correction. Data matrices were first converted to anndata objects where each dataset was annotated under a new observation column “sample”, followed by merging (concatenate command). Poor quality cells were removed by filtering out cells (scanpy.pp.filter_cells() command) based on number of counts and mitochondrial genes percentage. Cells with fewer than 200 genes and with higher than 0.5% mitochondrial genes were excluded. Genes expressed in less than three cells were considered outliers and were excluded by filtering in the pre-processing step. To allow counts to become comparable among all cells, library-size was corrected by normalizing (scanpy.pp.normalize_total() command) to a scale factor of 10,000, followed by log-transformation (sc.pp.log1p() command) and annotation of highly variable genes (scanpy.pp.highly_variable_genes() command) with n_top_genes set to None. Scaling of the data to follow a linear-regression model (scanpy.pp.scale() command) was performed with max_value of 10. Dimensionality reduction was computed using linear principal component analysis (scanpy.tl.pca() command) and number of significant principal components (PC) to be used for clustering were determined by plotting each PC’s variance ratio. Next, selected PCs were used to calculate neighbourhood graph (scanpy.pp.neighbors() command) that is automatically embedded to compute uniform manifold approximation and projection ^61^ (UMAP) graph (scanpy.tl.umap() command) for two dimensional visualization of cells. Communities of cells were detected by Leiden ^62^ algorithm (scanpy.tl.leiden() command) with the default resolution of 1. ComBat algorithm was then used (scanpy.pp.combat() command) to remove batch variation between samples, and significant PC’s were selected to calculate neighbourhood graph and UMAP topology.

### Human sample gene matrix pre-processing, filtering, and cell clustering

Doublets in human sample matrix were high due to not using FACS. Doublets were removed from raw count matrix prior to pre-processing by Scrublet ^63^ (ver. 0.2.3) python package with upstream data log transformation. Scrublet object was initialized (scrublet.scrublet() command) with sparse raw count matrix and expected doublet rate of 0.076. Scores were calculated and binary prediction was assigned for each cell (scrublet.scrub_doublets() command) with 30 PCs and log_transform set to True. Cells predicted as True doublets were removed from the count matrix. Scanpy was used for further pre-processing, filtering, and clustering of doublet-free matrix. Analysis was performed as described for mouse without batch correction and with few modifications. Cells with fewer than 500 genes and genes expressed in less than ten cells were considered outliers and were excluded by filtering. Annotation of highly variable genes (scanpy.pp.highly_variable_genes() command) with n_top_genes is set to 2000 and with flavor set to Seurat_3. To completely exclude doublets prior to cardiomyocytes sub-clustering, cells with high expression of one other cell marker gene were removed, based on expression levels and frequency, we removed cells expressing *VWF*. For the integration with healthy sample, single nuclei Robj.gz file of 11 years donor sample ^36^ was downloaded from the human cell atlas database (https://data.humancellatlas.org/explore/projects/135f7f5c-4a85-4bcf-9f7c-4f035ff1e428). Robj file was converted from R object to h5ad file in R (ver. 4.1.0) using sceasy ^64^ (ver. 0.0.6) library (sceasy::convertFormt command) then used in Scanpy for downstream integration. BBKNN ^65^ (ver. 1.5.1) integration was then performed on non-scaled, logtransformed, and normalized concatenated anndata object (scanpy.external.pp.bbknn command) using sample as the batch key including 30 PCs. UMAP was finally run on the integrated space (scanpy.tl.umap command).

### Differential gene expression analysis and annotations

To identify highly expressed genes in Leiden clusters, differentially expressed genes were ranked (scanpy.tl.rank_genes_groups() command) using Wilcoxon rank-sum test. Genes were ranked by log_2_ fold change (log_2_FC) values and *p*-values. Top statistically significant genes were used for Cell type manual annotation based on the expression of cell-specific marker genes manually curated from literature.

### Sub-clustering

Cardiomyocytes were isolated and re-clustered, UMAP graph was computed, and Leiden algorithm was used with 1 resolution. For the sake of reproducibility, sub-clustered cardiomyocytes were stored as anndata file and further analyzed. Differentially expressed genes analysis between different samples was then performed using Wilcoxon rank-sum test and genes were ranked by log_2_FC values and *p*-values. Ranked genes lists were used for comparative analysis and downstream gene ontology analysis.

### Pseudo-time trajectory analysis

Integrated re-clustered cardiomyocytes were used for trajectory analysis. ForceAtlas2 ^66^ (FA) (ver. 0.3.5) algorithm was used to draw single-cell graph (scanpy.tl.draw_graph() command) using the anndata object previously calculated PCA space and neighbourhood graph. To denoise the graph, diffusion maps were computed (scanpy.tl.diffmap() command) using the first 15 components and neighbourhood distances were calculated between diffusion components (scanpy.pp.neighbors() command) to draw graph on the diffusion map (scanpy.tl.draw_graph() command). Clusters were detected by Leiden algorithm (scanpy.tl.leiden() command) with the default resolution of 1. Next, PAGA^29^ (ver. 1.2) graph was calculated (scanpy.tl.paga() command) based on embeddings connectivity maps of UMAP clusters calculated by the Leiden algorithm. The single-cell graph was recomputed (scanpy.pl.draw_graph() command) based on PAGA graph calculated earlier. To assign a root for pseudotime temporal path tracing, the first cluster with the highest *Ppargc1a* expression was chosen.

### Gene ontology analysis

Gene lists were filtered with a *p*-value cut-off of <0.01 and log_2_FC value of >1.5. GO analysis was performed on gene lists using EnrichR ^67–69^ and *p*-values were calculated with Fisher exact test.

### Immunohistology staining

Freshly dissected heart tissues were fixed in ice-cold 4% Paraformaldehyde Phosphate Buffer Solution (4% PFA, WAKO) for 24 h followed by serial immersion of the tissue in 10%, 20%, 30% sucrose in PBS for 24 h each. Tissues were embedded in OCT compound (Sakura Finetek) and frozen by chilling at −80°C. For antigen retrieval, sections (8 μm) were submerged in 1X Citrate buffer, pH 6.0 (Abcam) and microwaved for 2 min followed by 10 min boiling in −98°C water bath. Sections cooled for at least 30 min at room temperature were submerged under running hot water for 1 min and washed with 1X buffer two times for 3 min. Sections were blocked with Blocking One Histo (Nacalai) for 10 min at room temperature, then incubated with primary antibodies at 4°C overnight. For double staining, 4°C overnight incubation of rabbit anti-Atf3 (1:500 Abcam Cat# ab254268) followed by three times PBS washing and then incubation with goat anti-cTnnI3 (1:500 Abcam Cat# ab56357) for an additional overnight at 4°C. Three times washing with PBS followed by secondary antibodies goat anti-rabbit IgG Alexa Fluor 555 (Abcam Cat# ab150078), donkey anti-goat IgG Alexa Fluor 488 (ThermoFisher Cat# A-11055), and Hoechst incubation for 1 h at room temperature Sections were washed and mounted with VECTASHIELD (VECTOR LABORATORIES). Images were obtained using confocal laser scanning microscope (Olympus, FV3000).

### Spatial transcriptomics

Spatial transcriptomics of Atf3-positive cells was performed using photo-isolation chemistry (PIC) as described ^32,33^. In brief, fresh-frozen tissue sections embedded in OCT compound (Sakura Finetek) after ice-cold PBS perfusion and frozen by incubating on isopentane chilled in liquid nitrogen were permeabilized with HCl (Nacalai Tesque), followed by *in situ* mRNA reverse transcription using photo-caged oligodeoxynucleotides (ODNs) primers (Glen research). After first strand synthesis, anti-Atf3 antibody was used to identify region of interest (ROI) by immunostaining. Tissue sections were divided into separate groups; high Atf3 and low Atf3 sections. UV irradiation (LED light, Prizmatix) was used for uncaging ODNs allowing in-vitro transcription reaction. Libraries were further reverse-transcribed and paired-end sequenced on NovaSeq6000 (Illumina). Final reads were mapped to reference genome (GRCm38) using HISAT2 ^70^ (ver. 2.1.0). Analysis was performed on the web-based integrated Differential Expression and Pathway ^71^ (ver. 1.1) Analysis (iDEP) tool. For DEG selection, two-tailed t test was performed on normalized processed counts extracted from iDEP, followed by log_2_FC calculations. Gene lists were filtered with a *p*-value cut-off of <0.05. GO analysis was performed on gene lists using EnrichR and *p*-values were calculated with Fisher exact test.

### RNA-sequencing

RNA was isolated from 12-week hearts of MyoAAV2-shCtrl or MyoAAV2-sh*Atf3* injected FS6KD mice using RNeasy fibrous kit (QIAGEN). Libraries were prepared for sequencing using a TruSeq Stranded mRNA sample prep kit (Illumina, San Diego, CA, USA) according to the manufacturer’s instructions. Whole transcriptome sequencing was performed using NovaSeq6000 platform (Illumina) in a 101 bp single-end mode. Sequenced reads were mapped to the mouse reference genome sequence (GRCm38_p6). BAM files were then each converted into one integrated count file using FeatureCounts (ver 2.0.0). Read count file was normalized by the CPM (counts per million) method using iDEP. Genes expressed with >1 CPM in at least one of the samples of any condition were only used for the analysis. For DEG selection, two-tailed t test was performed on normalized processed counts extracted from iDEP, followed by log_2_FC calculations. Gene lists were filtered with a *p*-value cut-off of <0.05. GO analysis was performed on gene lists using EnrichR and *p*-values were calculated with Fisher exact test.

### Statistical analysis

Data were analyzed using GraphPad Prism 8 and were considered significant when *P* < 0.05. Data between two groups were compared with Student’s *t*-test. Data from more than two groups were evaluated with ordinary one-way ANOVA followed by Tukey’s *post hoc* analysis, and data from compound two or more groups were evaluated with two-way ANOVA followed by Tukey’s *post hoc* analysis. All data are presented as mean + SD. Schematic diagrams were created and images were processed by Adobe Illustrator (ver. 27.0.1).

