## Supplemental figures for "Atf3 controls transitioning in female mitochondrial cardiomyopathy as identified by single-cell transcriptomics"

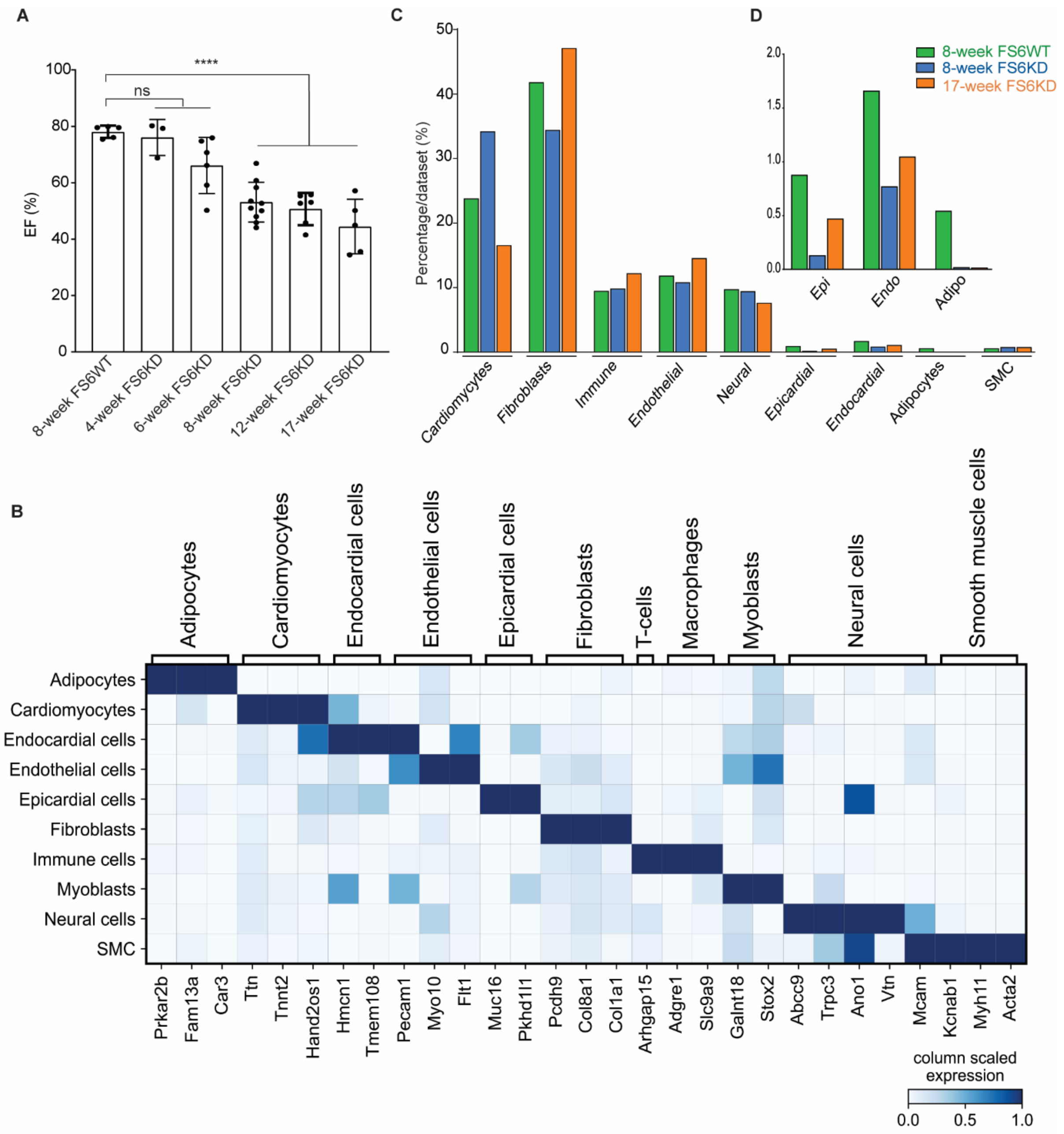


**Figure S1. Female FS6KD mice demonstrated progressing phenotype and different cellular composition.**

(A) Serial EF measurements of female FS6KD mice starting 4 weeks in comparison to 8-week FS6WT mice. Bars: mean ± SD. Dots: individual subjects. Ordinary one-way ANOVA and Tukey’s for multiple comparisons test for age comparisons, statistical significance: ***p ≤ 0.001, ns; not significant.

(B) Heatmap displaying expression of different cardiac cellular populations marker genes generated from the integrated dataset and their cell type annotations.

(C and D) Percentages of cellular populations per sample within the integrated dataset with (D) focus on percentage of epicardial cells, endocardial cells, and adipocytes.

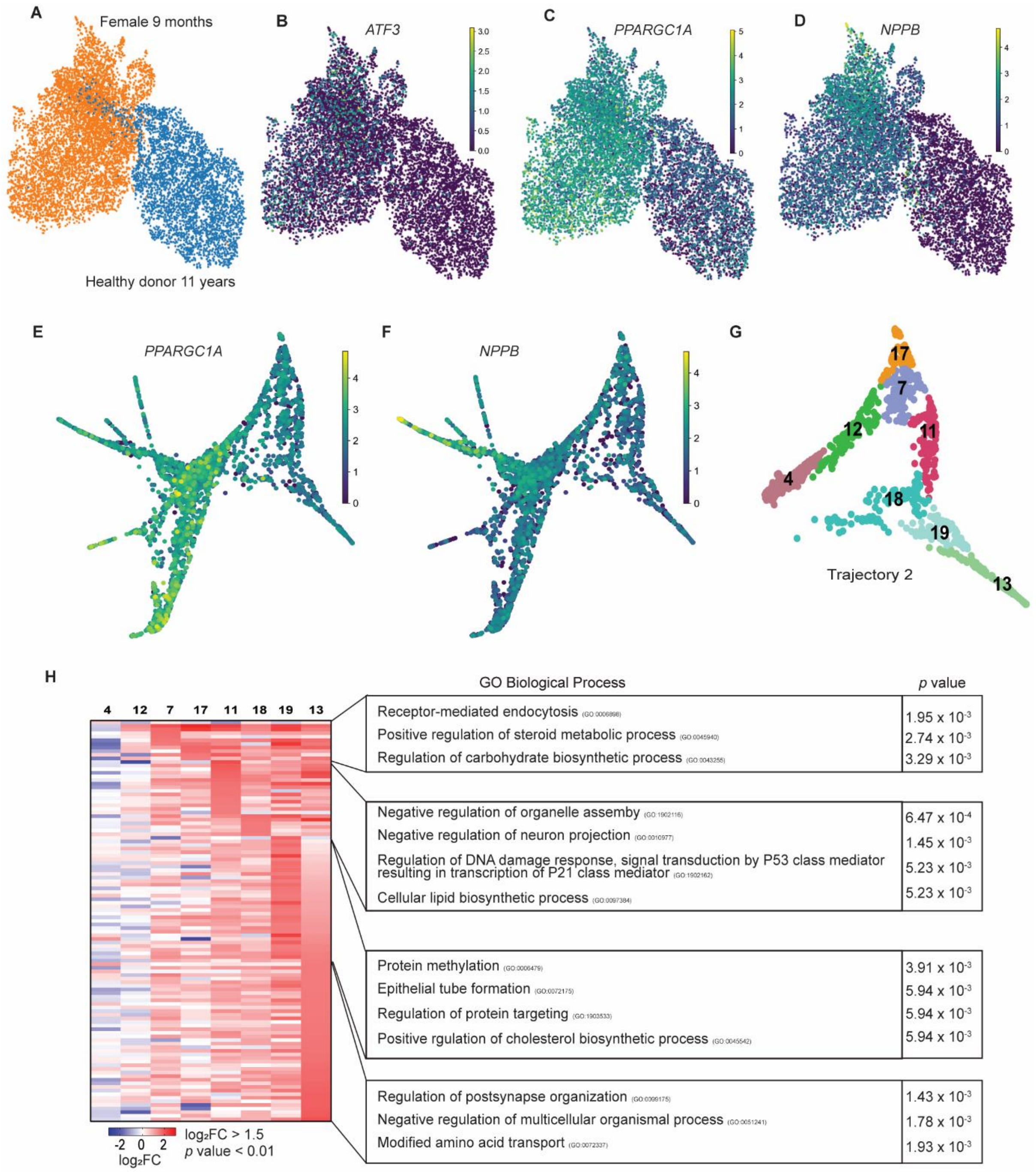


**Figure S2. Human sample showed more cardiomyocytes heterogeneity compared to mice.**

(A) UMAP plot of cardiomyocytes from our human MD patient integrated with ventricular cardiomyocytes isolated from 11 year old female donor dataset colored by sample identity, our sample (orange) and donor sample (blue). Expression shown as cardiomyocytes UMAP features of (B) *ATF3,* (C) *PPARGC1A*, and (D) *NPPB.*

(E) *PPARGC1A* expression shown as FA2 scatter plot.

(F) *NPPB* expression shown as FA2 scatter plot

(G) FA2 scatter plot of PAGA trajectory from cardiomyocytes numbered by leiden cellular states, showing only the second trajectory.

(H) Heatmap displaying top upregulated genes in the cellular states of the second trajectory selected cellular states ranked by $\log_2$FC and associated top GO terms. Fisher exact test was used to calculate *p*-values for GO.

Related to Main Figure 6
